# Optimal Random Avoidance Strategy in Prey-Predator Interactions

**DOI:** 10.1101/2020.03.04.976076

**Authors:** Masato S. Abe, Minoru Kasada

## Abstract

It has recently been reported that individual animals, ranging from insects to birds and mammals, exhibit a special class of random walks, known as Lévy walks, which can lead to higher search efficiency than normal random walks. However, the role of randomness or unpredictability in animal movements is not very well understood. In the present study, we used a theoretical framework to explore the advantage of Lévy walks in terms of avoidance behaviour in prey-predator interactions and analysed the conditions for maximising the prey’s survival rate. We showed that there is a trade-off relationship between the predictability of the prey’s movement and the length of time of exposure to predation risk, suggesting that it is difficult for prey to decrease both parameters in order to survive. Then, we demonstrated that the optimal degree of randomness in avoidance behaviour could change depending on the predator’s ability. In particular, Lévy walks resulted in higher survival rates than normal random walks and straight movements when the physical ability of the predators was high. This indicates that the advantage of Lévy walks may also be present in random avoidance behaviour and provides new insights into why Lévy walks can evolve in terms of randomness.

## 1. Introduction

To an observer, animal behaviour often seems random and unpredictable. Even when individual animals are subjected to the same experimental apparatus or external stimuli, behavioural variability is often observed [1–5]. It has been suggested that, to some extent, such variability stems from the nervous system that can initiate spontaneous actions [1,3,5–8]. However, how is variability or randomness in animal behaviour associated with evolutionary advantages? One of the candidate hypotheses is that such variability may improve a prey’s efficiency in avoiding a predator’s attacks in prey-predator interactions [6,9,10]. If the prey moves in the same manner every time, the predator can easily predict its movement and predation will be successful. Apart from top predators in a food web, most organisms are exposed to predation risk; therefore, the adaptive evolution of strategies that cannot be easily predicted must occur in many taxa. Consequently, in order to gain a full understanding of the role of variability or randomness in animal movements, it is crucial to explore the kind of movements exhibited by prey under predation risk and what the optimal avoidance strategy is, based on the cost and benefit involved [11,12].

Studies on the escape responses of prey to predators’ attacks have examined this topic from the perspectives of behavioural ecology and neurophysiology [9,10,13–15]. In particular, animal escapology has focused on how animals escape from threatening stimuli in an extremely short time-scale (e.g. 5-10 ms) [10,16–18]. Empirical evidence has shown that the variability in the direction of the prey’s movement may be an adaptive trait as a response to the predator that approaches it [2]. In contrast, another important consideration in prey-predator interactions is that when the predator is far from a prey individual, it can predict its movement by employing a kind of interception strategy and then it attacks it for a longer time-scale. For instance, a bird flying in the sky predicts the movement of an insect walking on the ground and then tries to attack it (figure 1a). Another example is an insect, such as a mosquito or a cockroach, which escapes from a human who is trying to catch it. In these situations, the predator aims for the prey target and tries to catch it (figure 1a). Although the speed with which the predator attacks is much faster than the speed with which the prey escapes, there would be a gap between an actual prey position and a predicted position before the attack. Besides, the prey would seek refuge if this would help it elude some predator attacks. In such cases, unpredictable movements such as random walks (figure 1b) would benefit the prey, although such movements would pose the risk of delayed arrival to the refuge. Therefore, it may be crucial for prey to exhibit movements with randomness rather than escape responses, such as reactive responses characterised by an extremely short time scale.

**Figure 1.**
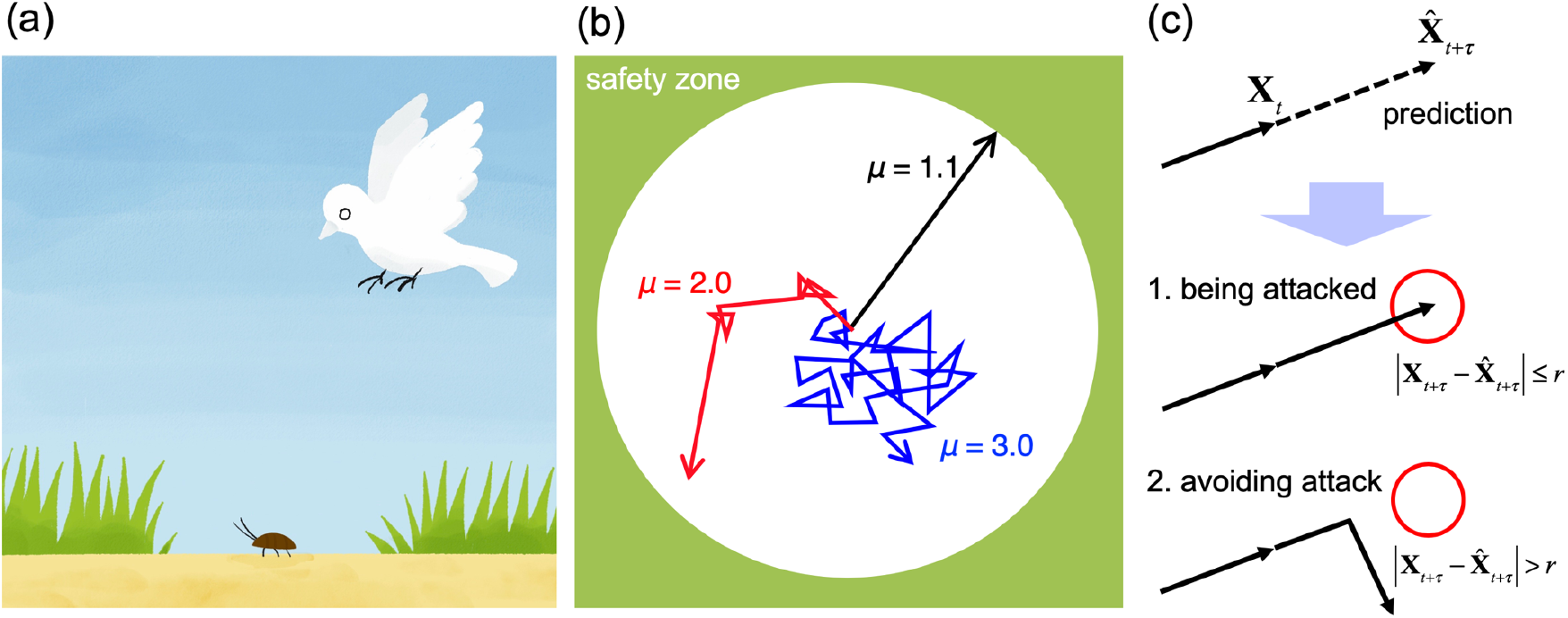
Random avoidance in prey-predator interactions. (a) We considered a scenario in which a predator observes from above a prey moving in an open area. (b) A predator’s view and the example of movement trajectories of a prey for *μ* = 1.1, 2.0 and 3.0. Note that *μ* = 1.1 and 3.0 roughly correspond to straight movement and Brownian walks, respectively. The green area is a safety zone. (c) How to predict the prey’s position. At time *t* the predator must predict the future (after time step *τ*) position of the prey. A red circle is an attack area with radius *r* and its centre is the predicted position 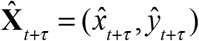. If the prey moves within the red circle after time step *τ*, the predator can be considered successful in having predicted the prey’s position and in attacking it. When the prey is outside the red circle after time step *τ*, the predator’s attack is regarded as a failure. Then, the predator tries to predict the next future position (after waiting time *W* has elapsed) and attack the prey. The solid arrows represent the actual trajectory of the prey’s movement and the dashed arrow represents the predator’s prediction.

It has recently been reported that various animals ranging from insects to fishes, birds, and mammals, including humans, exhibit a special class of random walks, called Lévy walks [19–25]. The step length *l* (i.e. the distance between consecutive reorientations) of Lévy walks follows a power law distribution, *P*(*l*)~*l^-μ^* where *μ*∈(1, 3] is a power law exponent characterising the movements. Lévy walks are characterised by rare ballistic movements among short steps (figure 1b), resulting in anomalous diffusions. Theoretical studies reported that Lévy walks have higher search efficiency than Brownian walks where the targets are distributed sparsely and patchily [26,27]. Therefore, it has been suggested that the prevalence of Lévy walks in animal movements is a consequence of natural selection that facilitates search efficiency. However, recent findings suggest that the various mechanisms that can produce Lévy walk patterns depend on several conditions [28,29]. Thus, it is imperative to explore why and when animals exhibit Lévy walks depending on various ecological conditions. Nevertheless, the impact of Lévy walks, other than search efficiency, on the fitness of individual animals remains poorly understood.

In the present study, we hypothesised that individual animals performing Lévy walks would benefit in terms of avoiding a predator’s attacks. The questions examined included what movement strategy is the most efficient in surviving in prey-predator interactions and how is the optimal behaviour affected by the ecological conditions and the predator’s cognitive and physical abilities [30]. To answer these questions, we constructed a general framework of the predator avoidance behaviour based on a random walk paradigm (figure 1a, b) and analysed the theoretical model to determine the efficiency of avoidance.

## 2. Methods

To explore the most efficient avoidance behaviour, we constructed a framework in which a prey individual avoided a predator that overlooked an open arena (figure 1a, b) and calculated the survival rate of the prey by using simulations and analytical solutions.

First, we considered a prey individual to be at the centre of the 2D circular field with radius *R* and a predator that overlooked the entire field from above (figure 1a, b). Then, the prey started moving and performed a class of random walks (i.e. Lévy walks with a specified *μ*) at a constant velocity (one space unit per unit time) without stopping. The step lengths *l* were drawn from a truncated power law distribution which is often best fitted to empirical data [19,20,23,24] and the turning angles between consecutive steps were drawn from a uniform distribution (see electronic supplementary material for details). In our analysis, *μ* ranged from 1.1 to 3.0 by 0.1. Lévy walks with *μ* = 1.1 and 3.0 correspond approximately to straight-line movements and Brownian movements, respectively.

We assumed that predators were rational and had relatively high cognitive ability. The predator could perceive the present position of the prey, denoted by **X**_*t*_ = (*x_t_*, *y_t_*), and could also memorise its past positions. Then, the predator would have to predict the future position **X**_*t*+*τ*_ = (*x*_*t*+*τ*_, *y*_*t*+*τ*_) of the prey after a *τ* time step, based only on the information it had about the present (*t*) and past positions of the prey. The predicted position was denoted by 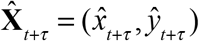. While the time lag *τ* could be considered as a predator attack characteristic, it could also be a characteristic of the prey individual as its position after *τ* could be scaled by its velocity. Therefore, if the prey’s velocity was fast, *τ* could be relatively large. Moreover, if the predator attacked slowly after making a prediction, *τ* could also become large. Intuitively, the larger the τ, the higher the prey’s advantage in escaping the attacks.

To model how the predator would predict the prey’s position in a simple manner, we assumed that the probability distributions of step lengths and turning angles of the prey did not depend on its spatial positions. Therefore, we could straightforwardly derive the optimal prediction rule adopted by predators by using a straight line (figure 1c). This is similar to interception strategies which have been observed in birds, bats, fishes, and insects [31–35]. The predicted position can be described as follows:

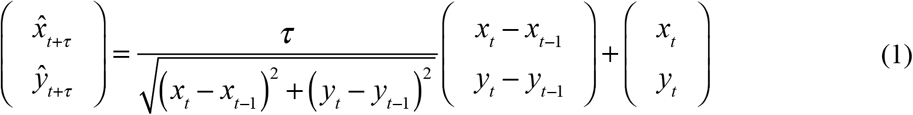

where (*x*_*t*-1_, *y*_*t*-1_) is the past position of the prey used to obtain the direction of the prediction (figure 1c). When the distance between the predicted position and the actual position was less than or equal to an attack radius *r*, the predator could successfully catch the prey. If the prediction failed, the predator would predict the new position again after the waiting time (*W*) had elapsed. *W* represents the time interval from the present attack to the next attack and is indicative of a characteristic of the predator that cannot attempt the next attack immediately. In contrast, once the prey reached the safety area (i.e. the edge of the circular field) (figure 1) without having been predicted by the predator, it would never leave it and would be regarded as having survived. To obtain the survival rate of the prey, this trial (from the beginning to the point of being caught or having survived) was independently iterated 10^6^ times for a specified parameter set in our simulation.

Although, based on our assumption, the prey could not obtain any information about when and where the predator would attack, we considered two cases: the prey was either aware (precaution case) or not aware (unwariness case) of the presence of the predator. The difference of these cases would result in a different step length at an initial condition (*t* = 0). In the precaution case, the prey could choose a new step length from the probability distribution at the initial condition and then it could start moving. On the other hand, in the unwariness case, the prey was on the move with step length l. Therefore, the initial step length is likely to be longer than that of the precaution case. Under these assumptions, we conducted simulations to estimate the survival rate of the prey.

To obtain an analytical solution of the survival rate based on the aforementioned assumptions, we split it into two components; the prediction rate and the time length to reach the safety area. In this case, the survival rate *ϕ* of a prey can be described as follows:

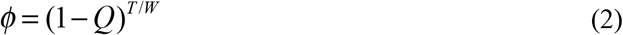

where *Q* is the predictability of the prey’s movements by the predators and *T*/*W* is the average number of attacks until the prey reaches the safety area. *Q* and *T* are functions of the parameter associated with the movements, namely *μ*, but the explicit derived expression can be obtained only for *Q* with *r* → 0 (see electronic supplementary material for details). Therefore, we used the simulations for *T*.

In our framework, the most unpredictable strategy included movements with many reorientations. In contrast, the straight-line movement was the easiest to predict. However, movements with reorientations could result in a delay in reaching the safety area, while straight-like movements could lead to short-term exposure to predation (figure 1c); the Lévy walk model could be an intermediate strategy between the two.

## 3. Results

First, we demonstrated the relationship between the predators’ prediction rate and the prey’s time length to reach the safety area for specified movement patterns, which were obtained from simulations and analytical solutions (see figure 2 for the precaution case and figure S1 for the unwariness case). The smaller the *μ* exponents, the shorter the time it would take prey to reach the safety area. This roughly corresponds to the mean squared displacement, which characterises the diffusion property and increases as the exponents decrease [19,26]. As for the prediction rate, when the exponents *μ* were small, it was easier for the predators to predict the prey’s position. This is because Lévy walks with large *μ* are characterised by frequent reorientations. Note that the prediction rate of Lévy walks with very small *μ* was close to one. This result indicates that there is a trade-off relationship between the predictability and the time to reach the safety area for any *τ* and *R*, that is, the shorter the time it takes to reach the safety area, the higher the predictability and vice versa. An intuitive explanation for these results is shown in figure 1b. The trade-off relationship also implies that the prey cannot decrease both parameters simultaneously, although both the lower predictability and the shorter time length to reach the safety area can lead to high survival rates for the prey. Therefore, we expect that Lévy walks with intermediate *μ* will represent a better strategy for increasing the survival rate compared with straight movements and Brownian walks (figure 2).

**Figure 2.**
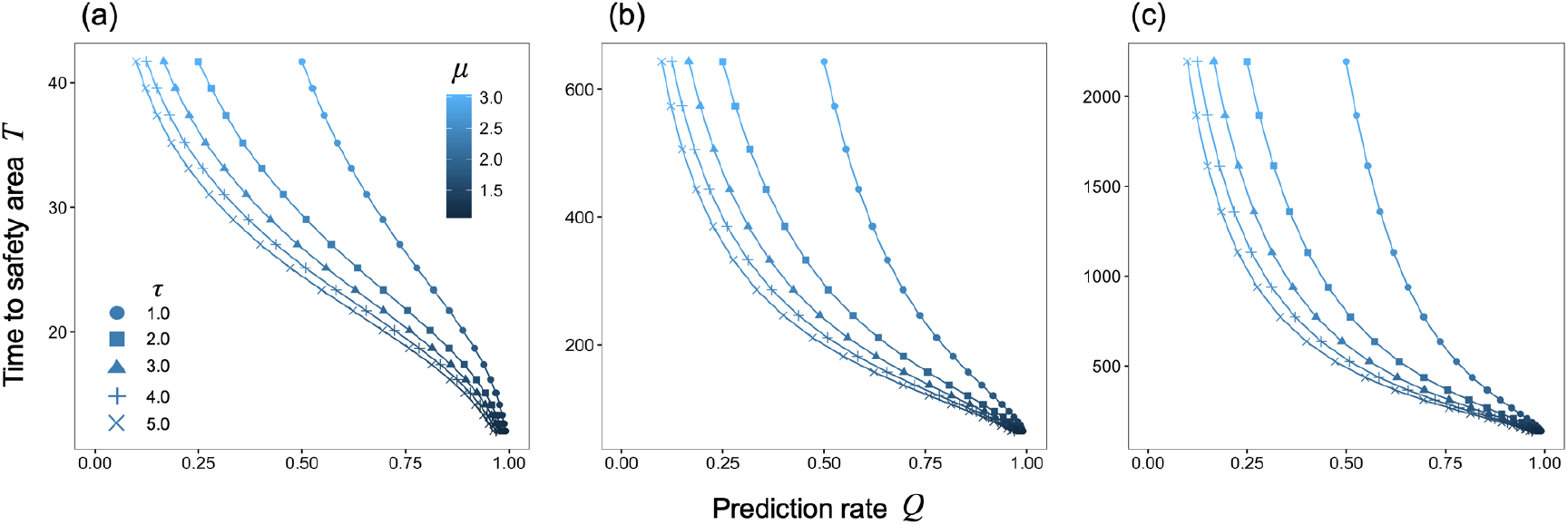
The relationship between the prediction rate and time length to reach the safety area. The horizontal axis is a prediction rate *Q* for *r* → 0 obtained by an analytical solution and the vertical axis is the average time length to reach the safety area for the precaution case, which was obtained by simulations with different area sizes (a) *R* = 10, (b) *R* = 50 and (c) *R* = 100. The colour represents the prey’s strategy parameter *μ*. The cut-off values of step length that we used were *l*_min_ = 1 and *l*_max_ = 1,000. The number of iterations for our simulation was 10^6^.

Next, we analysed the relationship between the survival rate and the movement pattern (*μ*) and examined the dependency of the other parameters (*τ, R*, and *W)* on this relationship. Figure 3 shows the results for the precaution case, suggesting that Lévy walks with intermediate *μ* can lead to a higher survival rate for *W* = 50 and 100, *R* = 50 and 100 and a small prediction time lag *τ*. In contrast, when *τ* was large, Brownian walks (i.e. Lévy walks with *μ* = 3) resulted in higher survival rates than Lévy walks with *μ* < 3 did in any scenario. When *τ* was small, the difference in the prediction rates for large and small *μ* was relatively small (figure 2). Therefore, the advantage of Brownian walks was negligible in terms of decreasing the prediction rate.

**Figure 3.**
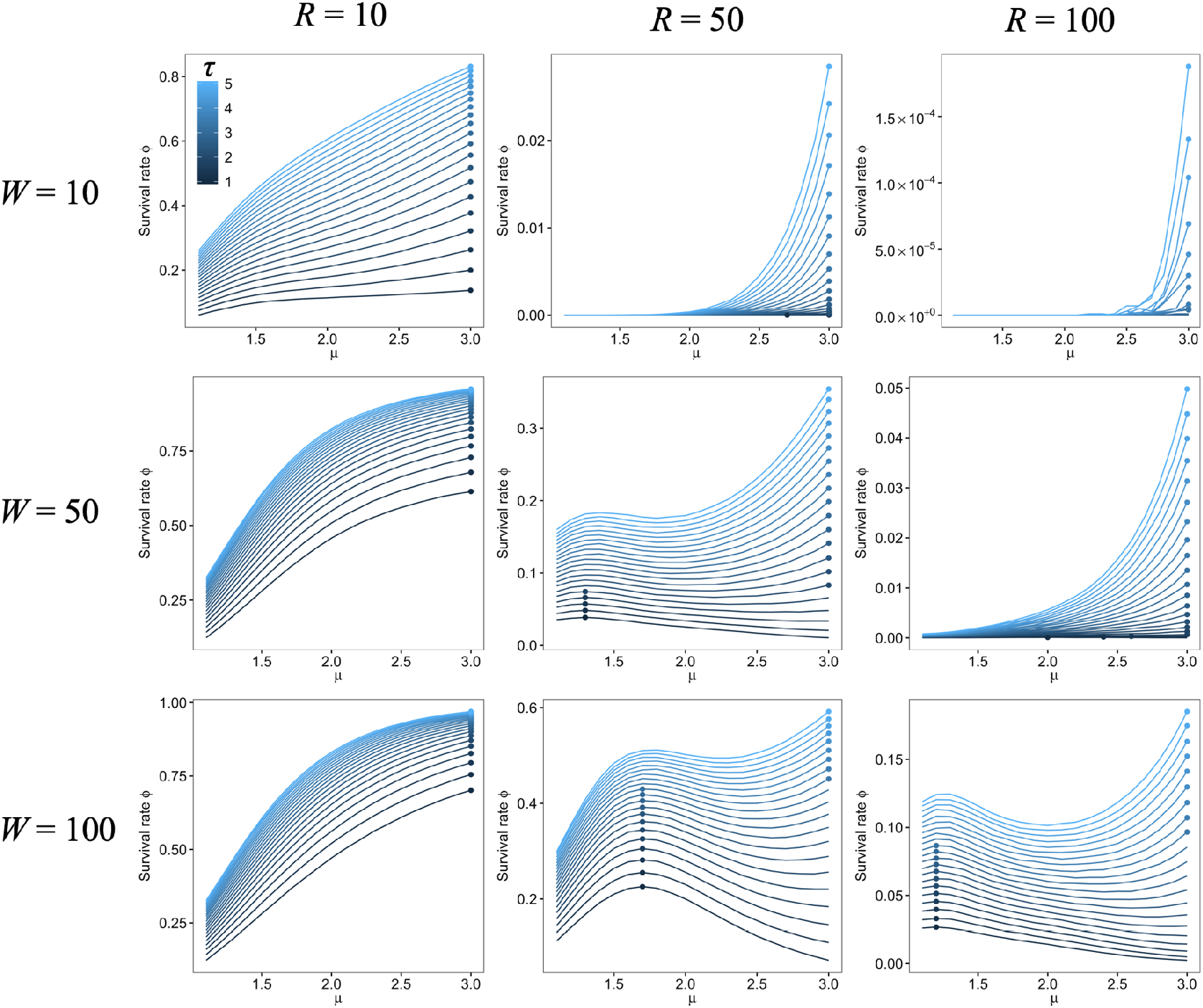
The simulated survival rate for the precaution case depended on μ. The horizontal axis is a prey’s strategy parameter *μ* and the vertical axis is the survival rate of the prey. The colour indicates the prediction time lag *τ*. The points represent the maximum survival rate for a specified parameter set (*R*, *W*, and *τ*). The cut-off values of step length that we used were *l*_min_ = 1 and *l*_max_ = 1,000. The attack radius *r* was set as 0.01. The number of iterations for our simulation was 10^6^.

The results also showed that Brownian walks were favoured when the radius *R* of the area was small (*R* = 10). This could be because of the short time length to reach the safety area (e.g. *T*≈40), even for Brownian walks, compared with the time length to reach the area for *R* = 50 and 100 (figure 2). Therefore, for small *R*, Brownian walks resulted in few attacks with a low prediction rate. In contrast, for large *R*, Brownian walks were more likely to be attacked, owing to the longer time it took to reach the safety area.

The waiting time *W* was also a key factor in determining the best avoidance strategy, for *R* = 50 and 100 (figure 3) in particular. *W* could affect the number of possible attacks because the average number of attacks was represented by the *T*/*W* ratio. When *R* and *μ* were large, *T* could also be larger. In this case, Brownian walks caused more predator attacks, thereby resulting in a low survival rate in spite of the low prediction rate.

These tendencies were also observed in the unwariness case and the semi-analytical results for *r* → 0 (figures S2, S3, and S4). Our results suggest a general tendency of Lévy walks to result in a higher survival rate for small *τ*; for large *τ*, the lower prediction rate strategy (i.e. *μ* = 3) was more efficient.

## 4. Discussion

In the present study, we created a theoretical framework to investigate random avoidance behaviour in terms of Lévy walks, which have been observed widely in biological movements [19–25]. We constructed a framework for prey-predator interactions based on the prey’s movements and the predator’s prediction (figure 1) and calculated the predator’s prediction rate, the time length to reach the safety area and the prey’s survival rate. We found that there is a trade-off relationship between the predator’s prediction rate and the prey’s time length to reach the safety area (figure 2). We can interpret the latter as the degree of exposure to predation risk. Thus, the trade-off corresponds to the relationship between the predictability and the degree of exposure to predation risk, suggesting that prey individuals cannot decrease both of these factors, although they need to. Consequently, they must identify a point of compromise in order to achieve higher survival rates.

Then, we showed that Lévy walks could outperform Brownian walks in terms of the survival rate when the time lag *τ* between the prediction and the attack was short and the area was not too small (figure 3). This implies that if the predator attacks the prey fast after predicting its position, Lévy walks can be the most efficient avoidance strategy. The attack characteristics associated with the time lag *τ* can be classified into two categories; the predator’s physical and cognitive abilities. The short time lag (small *τ*) is equivalent to the high physical ability that results in fast attacks or in highly flexible changes of attack directions. Likewise, it can be suggested that if making a prediction is a, somewhat, time-consuming task, the predator’s high cognitive ability results in a shorter time lag; however, in our model we did not make explicit assumptions about time regarding the predator’s predictions. Moreover, as the prey’s slow movement corresponded to a small time lag *τ*, prey individuals with low physical ability may experience higher survival rates by employing Lévy walks. As the small size of the area may correspond to a situation where obstacles such as grass where a prey can immediately hide under are present, Lévy walks can be favoured in relatively large open areas. Thus, these results show that there is a crucial connection between the prey’s fitness and the ecological and physical conditions that result from the prey’s movements and the predator’s predictions.

When the time lag was large, Brownian walks represented a better escape strategy (figure 3). If the movements were sufficiently random, they decreased the chances of the prey’s position being predicted by the predator. Furthermore, when *W* was small, the predator had a higher chance of attacking; in this case, the random movements represented an advantage. In both cases, eluding the attack was relatively more important for the prey’s survival than reaching the refuge rapidly was. This result suggests that Brownian walks still represent an effective strategy of escaping from a predator, which aims for and catches prey.

So far, the main advantage of Lévy walks has been the high encounter rate with targets (e.g. food, mates and habitats), which are distributed scarcely and patchily [26,27]. However, recent findings suggest that the origins of Lévy walks may not be unique, but may depend on ecological and physical conditions [28,29]. Therefore, it is crucial to explore when and why animal individuals would exhibit Lévy walks and which ecological traits, physical constraints or environmental factors are responsible for this phenomenon. In this study, we focused on the prey-predator interactions once they encounter each other. This scenario occurs in relatively shorter time and smaller spatial scales than the scenario in which a predator moves in search for prey moving in the environment [36,37]. Our results suggest a novel advantage of Lévy walks in the context of avoiding a predator’s attacks. Moreover, although the essential role of variability or randomness in Lévy walks is not very well understood, our results indicate that the extent of randomness plays a key role in determining the optimal avoidance strategy. Additionally, based on our framework, Lévy walks should be expressed as spontaneous behaviour, which is generated from the internal state of the prey, as the prey determines the reorientation and step length independent of external factors such as environmental cues. Therefore, these results point to one of the evolutionary origins of spontaneously produced Lévy walks [38].

A hunting strategy by predators with high cognitive abilities would not just be a reflective response to their prey’s movements. It has been shown that predators, including vertebrates and invertebrates, can predict the future position of a prey by using internal models [31,39,40]. In our model, we assumed that predators simply predicted the prey’s future position based on the prey’s direction, which reflects the prediction that is based on internal models. The evolution of the neural circuits associated with this prediction may be driven by the movements of prey. Further studies could reveal the co-evolution of prey’s movements and predators’ cognitive abilities.

In our framework, the environmental conditions were limited (e.g. circle). Indeed, in natural conditions, complex obstacles may be present, however the trade-off between the prediction rate and the time of exposure to the predation risk would still exist. It is important that future research addresses how environmental heterogeneity and the strategy’s dependency on environmental influences, affect prey-predator interactions [41]. Furthermore, considering situations in which a predator and a prey predict one another’s movements, is important in helping us gain a full understanding of prey-predator interactions.

## Acknowledgements

We would like to thank Saeka Shoda-Abe for drawing the illustration.

## Authors’ contribution

M.S.A designed the study. M.S.A and M.K. conducted the analysis. M.S.A. wrote the first draft of the manuscript. All authors contributed to the final version of the manuscript, gave final approval for publication, and agree to be held accountable for the work performed therein.

## References

1. Maye A, Hsieh C, Sugihara G. 2007 Order in spontaneous behavior. PLoS ONE, e443.(doi:10.1371/journal.pone.0000443)

2. Domenici P, Booth D, Blagburn JM, Bacon JP. 2008 Cockroaches keep predators guessing by using preferred escape trajectories. Current Biology 18, 1792–1796. (doi:10.1016/j.cub.2008.09.062)

3. Proekt A, Banavar JR, Maritan A, Pfaff DW. 2012 Scale invariance in the dynamics of spontaneous behavior. Proceedings of the National Academy of Sciences 109, 10564–10569. (doi:10.1073/pnas.1206894109)

4. Wearmouth VJ, McHugh MJ, Humphries NE, Naegelen A, Ahmed MZ, Southall EJ, Reynolds AM, Sims DW. 2014 Scaling laws of ambush predator ‘waiting’ behaviour are tuned to a common ecology. Proc. R. Soc. B 281, 20132997. (doi:10.1098/rspb.2013.2997)

5. Briggman KL. 2005 Optical imaging of neuronal populations during decision-making. Science 307, 896–901. (doi:10.1126/science.1103736)

6. Brembs B. 2011 Towards a scientific concept of free will as a biological trait: spontaneous actions and decision-making in invertebrates. Proc. R. Soc. B 278, 930–939. (doi:10.1098/rspb.2010.2325)

7. Sorribes A, Armendariz BG, Lopez-Pigozzi D, Murga C, de Polavieja GG. 2011 The origin of behavioral bursts in decision-making circuitry. PLoS Comput Biol 7, e1002075. (doi:10.1371/journal.pcbi.1002075)

8. de Ruyter van Steveninck RR. 1997 Reproducibility and variability in neural spike trains. Science 275, 1805–1808. (doi:10.1126/science.275.5307.1805)

9. Humphries DA, Driver PM. 1970 Protean defence by prey animals. Oecologia 5, 285–302. (doi:10.1007/BF00815496)

10. Domenici P, Blagburn JM, Bacon JP. 2011 Animal escapology I: theoretical issues and emerging trends in escape trajectories. Journal of Experimental Biology 214, 2463–2473. (doi:10.1242/jeb.029652)

11. Davies NB, Krebs JR, West SA. 2012 An introduction to behavioural ecology. Hoboken, New Jersey: John Wiley & Sons.

12. Blumstein DT, Samia DS, Cooper WE. 2016 Escape behavior: dynamic decisions and a growing consensus. Current Opinion in Behavioral Sciences 12, 24–29. (doi:10.1016/j.cobeha.2016.08.006)

13. Edmunds M. 1974 Defence in animals: a survey of anti-predator defences. Harlow: Longman.

14. Weihs D, Webb PW. 1984 Optimal avoidance and evasion tactics in predator-prey interactions. Journal of Theoretical Biology 106, 189–206. (doi:10.1016/0022-5193(84)90019-5)

15. Evans DA, Stempel AV, Vale R, Branco T. 2019 Cognitive control of escape behaviour. Trends in Cognitive Sciences 23, 334–348. (doi:10.1016/j.tics.2019.01.012)

16. Domenici P, Blagburn JM, Bacon JP. 2011 Animal escapology II: escape trajectory case studies. Journal of Experimental Biology 214, 2474–2494. (doi:10.1242/jeb.053801)

17. Card G, Dickinson MH. 2008 Visually mediated motor planning in the escape response of Drosophila. Current Biology 18, 1300–1307. (doi:10.1016/j.cub.2008.07.094)

18. Dewell RB, Gabbiani F. 2012 Escape behavior: linking neural computation to action. Current Biology 22, R152–R153. (doi:10.1016/j.cub.2012.01.034)

19. Viswanathan GM, Da Luz MG, Raposo EP, Stanley HE. 2011 The physics of foraging: an introduction to random searches and biological encounters. Cambridge: Cambridge University Press.

20. Humphries NE, Weimerskirch H, Queiroz N, Southall EJ, Sims DW. 2012 Foraging success of biological Lévy flights recorded in situ. Proceedings of the National Academy of Sciences 109, 7169–7174. (doi:10.1073/pnas.1121201109)

21. Raichlen DA, Wood BM, Gordon AD, Mabulla AZP, Marlowe FW, Pontzer H. 2014 Evidence of Lévy walk foraging patterns in human hunter-gatherers. Proceedings of the National Academy of Sciences 111, 728–733. (doi:10.1073/pnas.1318616111)

22. Reynolds AM, Frye MA. 2007 Free-flight odor tracking in drosophila is consistent with an optimal intermittent scale-free search. PLoS ONE 2, e354. (doi:10.1371/journal.pone.0000354)

23. Humphries NE et al. 2010 Environmental context explains Lévy and Brownian movement patterns of marine predators. Nature 465, 1066–1069. (doi:10.1038/nature09116)

24. Sims DW, Humphries NE, Hu N, Medan V, Berni J. 2019 Optimal searching behaviour generated intrinsically by the central pattern generator for locomotion. eLife 8, e50316. (doi:10.7554/eLife.50316)

25. Nagaya N, Mizumoto N, Abe MS, Dobata S, Sato R, Fujisawa R. 2017 Anomalous diffusion on the servosphere: A potential tool for detecting inherent organismal movement patterns. PLoS ONE 12, e0177480. (doi:10.1371/journal.pone.0177480)

26. Bartumeus F, da Luz MGE, Viswanathan GM, Catalan J. 2005 Animal search strategies: a quantitative random-walk analysis. Ecology 86, 3078–3087. (doi:10.1890/04-1806)

27. Viswanathan GM, Buldyrev SV, Havlin S, da Luz MGE, Raposo EP, Stanley HE. 1999 Optimizing the success of random searches. Nature 401, 911–914. (doi:10.1038/44831)

28. Reynolds A. 2015 Liberating Lévy walk research from the shackles of optimal foraging. Physics of Life Reviews 14, 59–83. (doi:10.1016/j.plrev.2015.03.002

29. Reynolds AM. 2018 Current status and future directions of Lévy walk research. Biology Open 7, bio030106. (doi:10.1242/bio.030106)

30. Wilson AM et al. 2018 Biomechanics of predator–prey arms race in lion, zebra, cheetah and impala. Nature 554, 183–188. (doi:10.1038/nature25479)

31. Mischiati M, Lin H-T, Herold P, Imler E, Olberg R, Leonardo A. 2015 Internal models direct dragonfly interception steering. Nature 517, 333–338. (doi:10.1038/nature14045)

32. Ghose K. 2006 Steering by hearing: a bat’s acoustic gaze Is linked to Its flight motor output by a delayed, adaptive linear law. Journal of Neuroscience 26, 1704–1710. (doi:10.1523/JNEUROSCI.4315-05.2006)

33. Fabian ST, Sumner ME, Wardill TJ, Rossoni S, Gonzalez-Bellido PT. 2018 Interception by two predatory fly species is explained by a proportional navigation feedback controller. J. R. Soc. Interface 15, 20180466. (doi:10.1098/rsif.2018.0466)

34. Brighton CH, Thomas ALR, Taylor GK. 2017 Terminal attack trajectories of peregrine falcons are described by the proportional navigation guidance law of missiles. Proc Natl Acad Sci USA 114, 13495–13500. (doi:10.1073/pnas.1714532114)

35. McHenry MJ, Johansen JL, Soto AP, Free BA, Paley DA, Liao JC. 2019 The pursuit strategy of predatory bluefish (Pomatomus saltatrix). Proc. R. Soc. B 286, 20182934. (doi:10.1098/rspb.2018.2934)

36. Bartumeus F, Catalan J, Fulco UL, Lyra ML, Viswanathan GM. 2002 Optimizing the encounter rate in biological interactions: Lévy versus Brownian strategies. Phys. Rev. Lett. 88, 097901. (doi:10.1103/PhysRevLett.88.097901)

37. Abe MS, Shimada M. 2015 Lévy walks suboptimal under predation risk. PLoS Comput Biol 11, e1004601. (doi:10.1371/journal.pcbi.1004601)

38. Kölzsch A, Alzate A, Bartumeus F, de Jager M, Weerman EJ, Hengeveld GM, Naguib M, Nolet BA, van de Koppel J. 2015 Experimental evidence for inherent Lévy search behaviour in foraging animals. Proc. R. Soc. B 282, 20150424. (doi:10.1098/rspb.2015.0424)

39. Kawato M. 1999 Internal models for motor control and trajectory planning. Current Opinion in Neurobiology 9, 718–727. (doi:10.1016/S0959-4388(99)00028-8)

40. Wolpert D, Ghahramani Z, Jordan M. 1995 An internal model for sensorimotor integration. Science 269, 1880–1882. (doi:10.1126/science.7569931)

41. Wilson AM, Lowe JC, Roskilly K, Hudson PE, Golabek KA, McNutt JW. 2013 Locomotion dynamics of hunting in wild cheetahs. Nature 498, 185–189. (doi:10.1038/nature12295)

