## Supplementary Material for "Optimal Random Avoidance Strategy in Prey-Predator Interactions"

### 1. Generating Lévy walks

Lévy walks are characterised by a power law distribution, $P(l) \sim l^{-\mu} (1 < \mu \le 3)$. However, recent empirical findings suggest that animal movements often follow a truncated power law distribution, that is, a power law distribution with an upper cut-off [1,2]. Therefore, in our simulations, we derived each step length $l$ following the truncated power law distribution with $\mu$ by using a uniform random number $u \in (0, 1)$ and the following equation:

$$l = l_{\min} \left\{ u \left[ 1 - \left( \frac{l_{\max}}{l_{\min}} \right)^{1-\mu} \right] + \left( \frac{l_{\max}}{l_{\min}} \right)^{1-\mu} \right\}^{1/(1-\mu)} \tag{S.1}$$

where $l_{\min}$ and $l_{\max}$ represent the minimum and maximum step length, respectively [3]. In our analysis, we used $l_{\min} = 1$ and $l_{\max} = 1{,}000$. The reason for this was that the order of magnitude in empirical studies on Lévy walks ranges from 1.5 to approximately 4, although there have been variations [2].

### 2. Analytical solution for the prediction rate

Here, we obtained analytical results for the prediction rate $Q$ in the case when the attack radius $r$ was close to 0. The step length $l$ of Lévy walks followed a truncated power law distribution with an exponent $\mu$:

$$P_1(l) = \frac{P(l)}{\int_{l_{\min}}^{l_{\max}} P(x)dx}. \tag{S.2}$$

When an animal individual was observed performing Lévy walks, the probability density distribution that the animal has selected $l$ was calculated by a weighted step length distribution as follows:

$$P_2(l) = \frac{lP_1(l)}{\int_{l_{\min}}^{l_{\max}} xP_1(x)dx}. \tag{S.3}$$

Moreover, the probability that the rest step length is longer than $m$ was:

$$\frac{l-m}{l}. \tag{S.4}$$

The probability that the predator could predict the prey's position after time step $\tau$ from observing the prey was equivalent to the probability that the rest step length was longer than $\tau$ because the prey's movement velocity was 1 (space unit)/(time unit). Thus, we obtained the probability:

$$\begin{aligned}\int_{\tau}^{l_{\max}} \frac{l-\tau}{l} P_2(l)dl &= \int_{\tau}^{l_{\max}} \frac{(l-\tau)P_1(l)}{\int_{l_{\min}}^{l_{\max}} xP_1(x)dx} dl \\ &= \int_{\tau}^{l_{\max}} \frac{(l-\tau)l^{-\mu}}{\int_{l_{\min}}^{l_{\max}} x^{(-\mu+1)}dx} dl.\end{aligned} \tag{S.5}$$

Finally, we obtained the explicit expression of the prediction rate,

$$Q\left(\tau,\mu,l_{\min},l_{\max}\right) = \frac{l_{\max}^{2-\mu} - \tau^{2-\mu}}{l_{\max}^{2-\mu} - l_{\min}^{2-\mu}} + \frac{\tau(2-\mu)\left(\tau^{1-\mu} - l_{\max}^{1-\mu}\right)}{(1-\mu)\left(l_{\max}^{2-\mu} - l_{\min}^{2-\mu}\right)} \tag{S.6}$$

when $\mu \neq 2$. For $\mu = 2$, we set $h = 2 - \mu$, then

$$\begin{aligned}\lim_{h\to 0} Q &= \lim_{h\to 0}\left( \frac{l_{\max}^{h} - \tau^{h}}{l_{\max}^{h} - l_{\min}^{h}} + \frac{\tau h\left(\tau^{h-1} - l_{\max}^{h-1}\right)}{(h-1)\left(l_{\max}^{h} - l_{\min}^{h}\right)} \right) \\ &= \lim_{h\to 0} \frac{l_{\max}^{h} - \tau^{h}}{h} \frac{h}{l_{\max}^{h} - l_{\min}^{h}} + \lim_{h\to 0} \frac{\tau\left(\tau^{h-1} - l_{\max}^{h-1}\right)}{(h-1)} \frac{h}{\left(l_{\max}^{h} - l_{\min}^{h}\right)} \\ &= \frac{\log l_{\max} - \log \tau}{\log l_{\max} - \log l_{\min}} + \frac{\tau - l_{\max}}{l_{\max}} \frac{1}{\log l_{\max} - \log l_{\min}}\end{aligned} \tag{S.7}$$

was obtained.

The time length to reach the safety area was calculated by using a simulation without the predation by predators, namely it was an absorbing condition as a boundary condition. The survival probability was obtained from the prediction rate and time using equation (2) shown in the main text.

### Figures

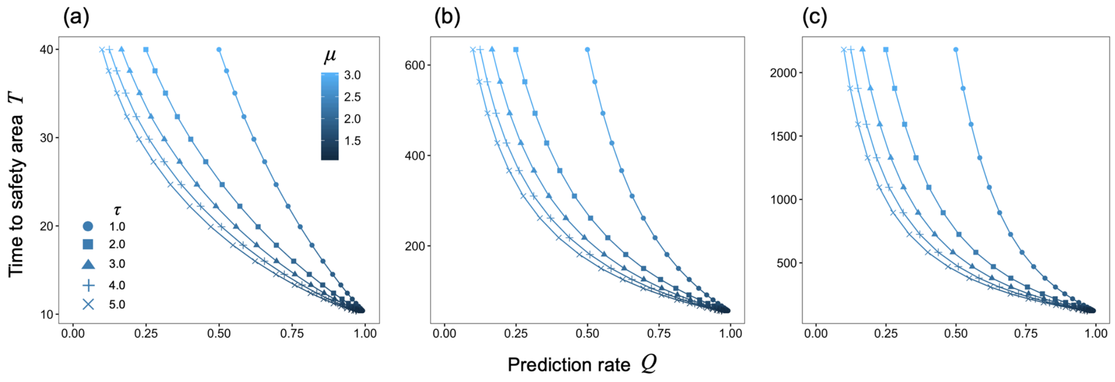


Fig. S1 The relationship between predictability and time of exposure to predation risk.

The horizontal axis is a prediction rate $Q$ for $r \rightarrow 0$ obtained by an analytical solution and the vertical axis is the average time to reach the safety area for the unwariness case obtained by simulations (a) $R$ = 10, (b) $R$ = 50 and (c) $R$ = 100. The colour represents the prey's strategy parameter $\mu$. The cut-off values of step length we used were $l_{\mathrm{min}} = 1$ and $l_{\mathrm{max}} = 1{,}000$. The number of iterations for our simulation was $10^6$.

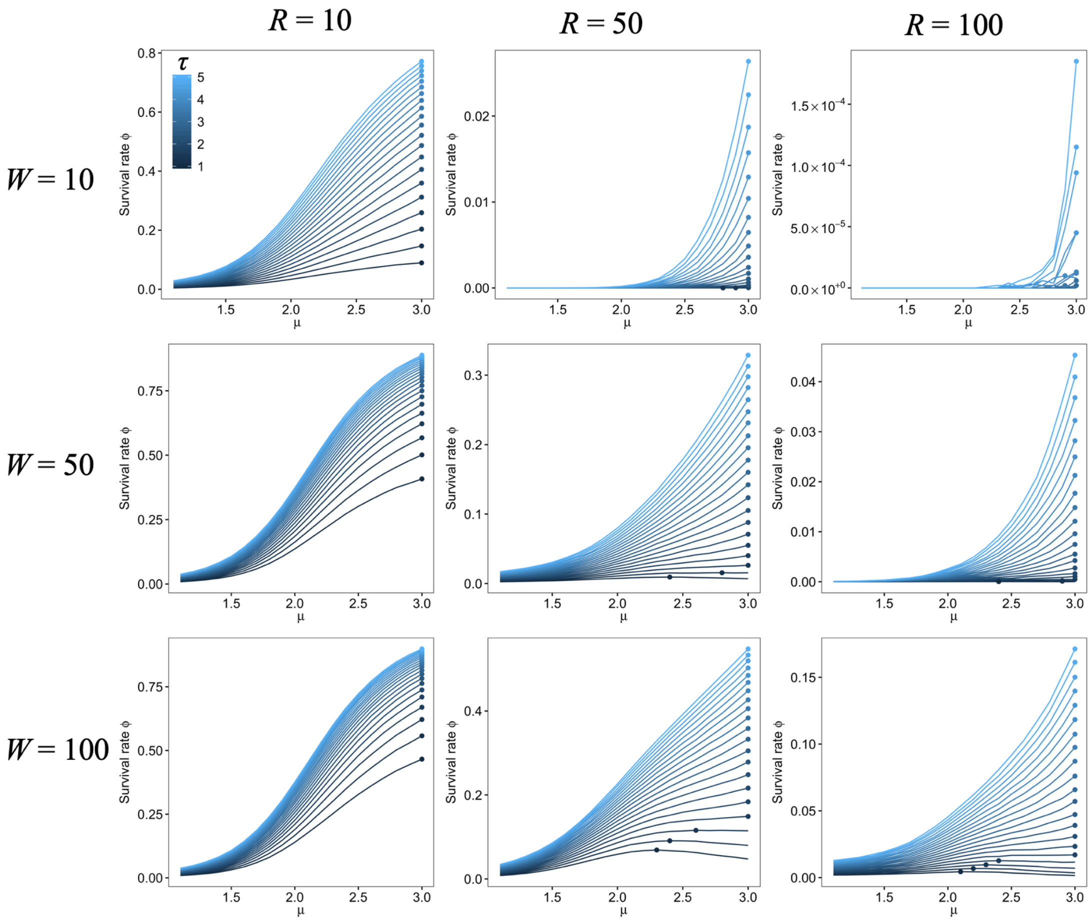


Fig. S2 The simulated survival rate for the unwariness case depended on $\mu$.

The horizontal axis is a prey's strategy parameter $\mu$ and the vertical axis is the prey's survival rate. The colour indicates the prediction time lag $\tau$. The points represent the maximum survival rate for a specified parameter set ($R$, $W$, and $\tau$). The cut-off values of step length we used were $l_{\mathrm{min}} = 1$ and $l_{\mathrm{max}} = 1{,}000$. The attack radius $r$ was set as 0.01. The number of iterations for our simulation was $10^6$.

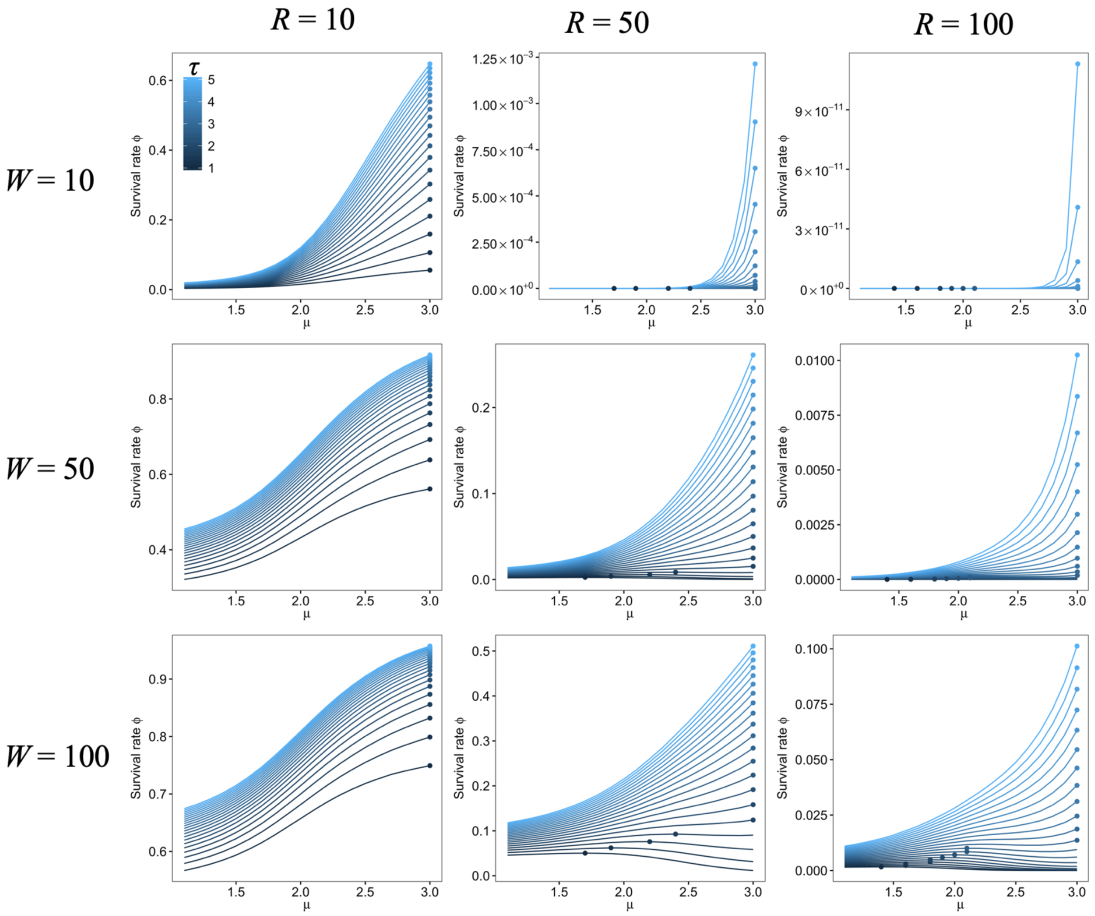


Fig. S3 The semi-analytical result of the survival rate for the precaution case depended on $\mu$. The horizontal axis is a prey's strategy parameter $\mu$ and the vertical axis is a prey's survival rate. The colour indicates the prediction time lag $\tau$. The points represent the maximum survival rate for a specified parameter set ($R$, $W$, and $\tau$). The cut-off values of step length we used were $l_{\min} = 1$ and $l_{\max} = 1{,}000$. The attack radius $r$ was set as 0.01. The number of iterations for our simulation was $10^6$.

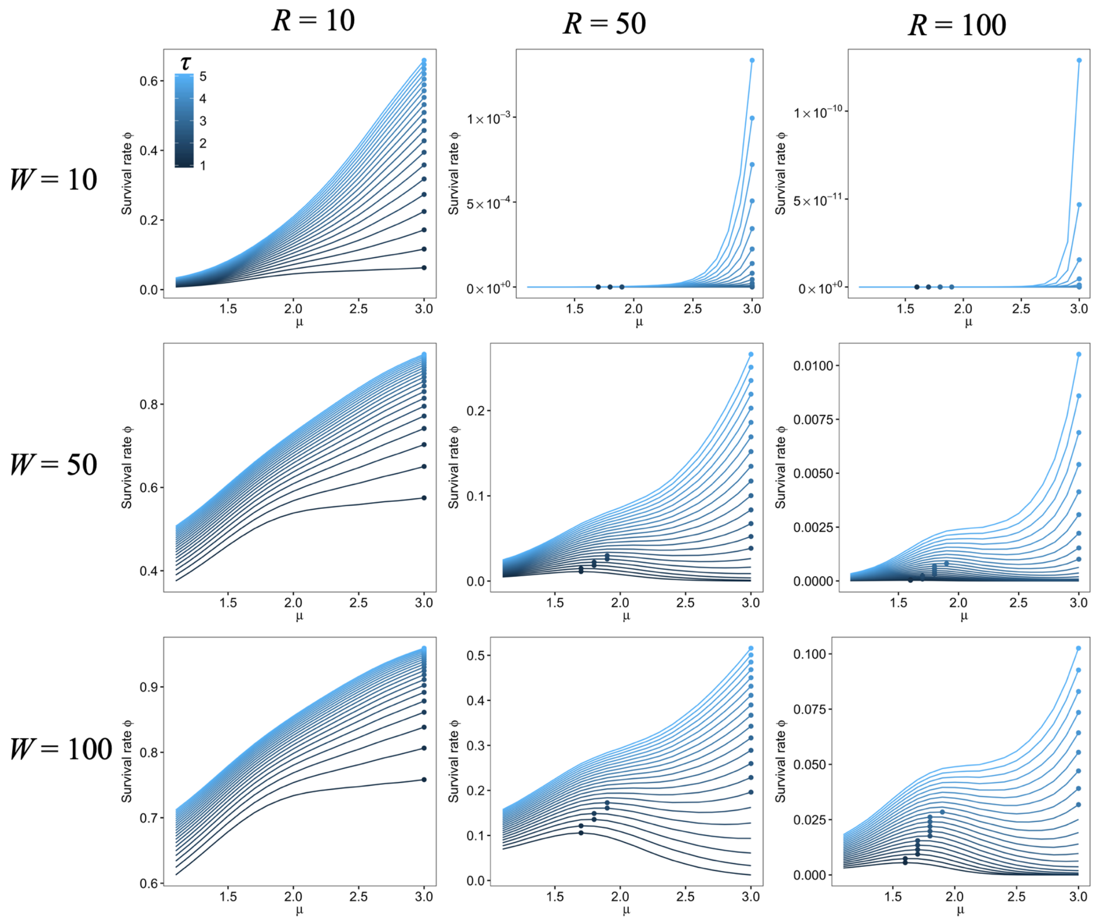


Fig. S4 The semi-analytical result of the survival rate for the unwariness case depended on $\mu$. The horizontal axis is a prey's strategy parameter $\mu$ and the vertical axis is a prey's survival rate. The colour indicates the prediction time lag $\tau$. The points represent the maximum survival rate for a specified parameter set ($R$, $W$, and $\tau$). The cut-off values of step length we used were $l_{\min} = 1$ and $l_{\max} = 1{,}000$. The attack radius $r$ was set as 0.01. The number of iterations for our simulation was $10^6$.

### References


1. Humphries NE *et al.* 2010 Environmental context explains Lévy and Brownian movement patterns of marine predators. *Nature* **465**, 1066–1069. (doi:10.1038/nature09116)

2. Humphries NE, Weimerskirch H, Sims DW. 2013 A new approach for objective identification of turns and steps in organism movement data relevant to random walk modelling. *Methods Ecol Evol* **4**, 930–938. (doi:10.1111/2041-210X.12096)

3. Kagan YY. 2002 Seismic moment distribution revisited: I. Statistical results: Seismic moment

distribution: I. *Geophysical Journal International* **148**, 520–541. (doi:10.1046/j.1365-246x.2002.01594.x)
